# Evaluating the Anti-aging Effects of Chinese Herbal Medicine *Rhodiola rosea* L. in Cultured Keratinocytes

**DOI:** 10.1101/2021.04.26.441423

**Authors:** Wu Han Toh, Chun-Bing Chen, Jude Clapper, Cheng-Chang Hsu, Hua-en Lee, Wen-Hung Chung

## Abstract

Among the many health benefits Chinese herbal medicine (CHM) presents, anti-aging is of special interest (Shen, Jiang, Yang, Wang, & Zhu, 2017; Zhao & Luo, 2017). Reported to possess anti-aging effects, the CHM *Rhodiola rosea* L., known colloquially as the “golden root”, has been widely incorporated in various drinks, daily supplements, and even cosmetics. This study investigates the effects of commercial *R. rosea* extracts and natural *R. rosea* roots on preventing UV-induced photoaging of the skin and correlates such effects with the composition of active ingredients in the extracts. To simulate the photoaging process, drug treated HaCaT cells were exposed to UVA and UVB radiation. The pharmacological anti-aging effects of *R. rosea* extracts were evaluated qualitatively and quantitatively through confocal immunofluorescence images with γ-H2A.X marker and telomerase activity assay (Telo TAGGG Telomerase PCR-ELISA). Preparatory thin layer chromatography and high-performance liquid chromatography were performed to isolate and quantify active ingredients. Cultured HaCaT cells showed morphological change after exposure to both UVA (>15.0 J) and UVB (>2000 mJ). The photoaging of keratinocytes was rescued by pretreating cells with *R. rosea* extracts as well as salidroside and rosavin active ingredients (P < 0.05). *R. rosea*-treated cells were characterized by increased telomerase activity and fewer γ-H2A.X foci compared to that of the control. Extracts with better preventative effects contained higher salidroside and rosavin content. The findings in this study reaffirm *R. rosea’s* efficacy as an anti-aging remedy and provide a basis for CHM’s integration into the mainstream of global healthcare.

## INTRODUCTION

Chinese herbal medicine (CHM) is a branch of traditional medicine based on over 3,500 years of medical practice. Alongside treatments like acupuncture and moxibustion, the practice is widely regarded to have distinct roots in Asia. However, in the past decade, CHM has gradually moved into the mainstream of global healthcare, emerging as a compliment, if not alternative to western medication (Lu & Lu, 2014). With its rising popularity, CHM’s many health benefits have garnered increased attention from both the general public and in scientific circles; of particular interest, are CHM’s purported anti-aging capabilities (Shen et al., 2017; Zhao & Luo, 2017).

Aging is an irreversible and inevitable process that affects all parts of the human body. On the cellular level, aging is associated with senescence, the cessation of cell division driven by mechanisms such as telomere shortening and other forms of genotoxic stress(Di Micco, Krizhanovsky, Baker, & d’Adda di Fagagna, 2021). Senescent cells are characterized by compromised nuclear structure, as well as enlarged and irregular shape. These changes are responsible for the progressive loss of physiological integrity observed on the organismal level (López-Otín, Blasco, Partridge, Serrano, & Kroemer, 2013). Although complete prevention of aging seems unlikely, anti-aging remedies that focus on delaying this process hold considerable pharmaceutical potential.

Known colloquially as the “golden root,” *Rhodiola rosea* L. is a Chinese herb that has been widely incorporated into various food additives, drinks, and cosmetic products. *R. rosea* plants are botanical adaptogens that grow mainly in the Himalayan belt and Tibet, and according to the basic tenets of traditional Chinese medicine, boost qi and reduce fatigue (Pu et al., 2020). In spite of the rich lore surrounding the herb, *R. rosea’s* pharmacological anti-aging effects have been well documented. Studies have verified *R. rosea*’s therapeutic value for the treatment of aging-related diseases, including Alzheimer’s disease, Parkinson’s disease, cerebrovascular disease, diabetes, and cardiovascular disease (Zhuang et al., 2019). In addition, in vitro studies performed in human lung fibroblast cell lines have demonstrated *R. rosea’s* ability to reverse senescence-like phenotypes (Chiang, Chen, Wu, Wu, & Wen, 2015). The root has also been shown to improve motor activity in mice and increase the lifespan of *Drosophila melanogaster* (common fruit fly) (Jafari et al., 2007; Mao et al., 2010). Current investigations of *R. rosea’s* biological activity focus mainly on its active components of salidroside, tyrosol, cinnamyl alcohol, rosavin, and rosarin (Choe et al., 2012; Dimpfel, Schombert, & Panossian, 2018; Kucinskaite et al., 2007).

This study evaluates *R. rosea’s* potential to retard aging of the skin, the largest and most accessible organ in the human body. While aging of internal organs are masked from the naked eye, the skin provides the first obvious marks of the passage of time, and therefore is often used to judge a person’s overall health. Rates of skin aging, while influenced in part by intrinsic genetic factors, can also be accelerated by exogenous environmental factors, ultraviolet (UV) irradiation chief among them (Ganceviciene, Liakou, Theodoridis, Makrantonaki, & Zouboulis, 2012). UV exposure from sunlight can be classified into UVC, UVB, and UVA wavelengths. UVC (100-290 nm) is largely obstructed by the ozone layer and has little impact on the skin. UVB (290-320 nm) penetrates into the epidermis and has strong mutagenic and carcinogenic effects on keratinocytes. UVA (320-400 nm) has deeper penetration and mediates chronic skin damage to both the epidermis and dermis. Although UVA is less energetic than UVB, it comprises a much larger portion of ambient light (Wang et al., 2019). Both UVB and UVA contribute to premature skin aging, a process known as photoaging (Gromkowska-Kępka, Puścion-Jakubik, Markiewicz-Żukowska, & Socha, 2021). Although past studies have posited general theories on the anti-aging properties of *R. rosea*, its effect on the skin is largely unknown. There are significant cosmetic applications if *R. rosea* is found to have preventative effects on photoaging.

Despite CHM’s growing popularity, concerns about commercialized CHM, *R. rosea* included, still inhibit its internationalization. For one, there is a lack of standardization of ingredients in commercial *R. rosea*, resulting in issues of quality control (Booker et al., 2016; Zhou, Li, Chang, & Bensoussan, 2019). Moreover, the quantification of the anti-aging effects of *R. rosea* herbal products remains unclear. The dose-effect relationship of the polyphenols contained in *R. rosea* has yet to be established, and the ways in which these compounds interact is unknown. Without sufficient knowledge on this front, self-administration of herbal products poses a significant health risk. The increased global consumption of *R. rosea* calls for more conclusive research regarding the root’s composition and its anti-aging effects. To date, this information is not available in most commercialized products (Lu & Lu, 2014).

The purpose of this study is to validate, quantify, and account for the anti-aging effects of *R. rosea* on the skin. As such, the present study consists of two main objectives. In the first objective, we investigate the anti-aging effects of *R. rosea* extracts on the human skin. The extent to which commercialized *R. rosea* delays UV induced cellular senescence of skin cells (i.e., keratinocytes) will be assessed through various aging biomarkers. In the second objective, we analyze the composition of active components salidroside and rosavin across different commercial *R. rosea* extracts. This information will reveal the ingredients predominantly responsible for *R. rosea’s* anti-aging effects, and provide a basis for CHM’s standardization, and eventually, modernization.

This study uses cellular morphology change, telomerase activity, and immunofluorescence staining of γ-H2A.X as biological markers of cell photoaging. Telomerase is a ribonucleoprotein DNA polymerase that maintains the length of telomeres, specialized repetitive DNA sequences found at the ends of linear chromosomes. In the absence of telomerase activity, telomeres progressively shorten due to the inability of DNA polymerase to replicate single-stranded 3’ ends (Bernadotte, Mikhelson, & Spivak, 2016; Hornsby, 2007). Cells that possess sufficient telomerase activity circumvent the replicative senescence or crisis states induced by telomere shortening (Nicholls, Li, Wang, & Liu, 2011). Telomerase has been found to play a vital role in the self-renewing skin, where it maintains skin function and proliferation (Buckingham & Klingelhutz, 2011). It has been suggested that telomerase activation by natural molecules can play a role in the treatment of aging-related diseases (Tsoukalas et al., 2019). In sum, as telomerase offsets cellular aging, its activity serves as a biological marker for cell health. TRAP (telomeric repeat amplification protocol) combined with colorimetric detection by ELISA (enzyme-linked immunosorbent assay) allows for the quantitative detection of telomerase activity in a highly efficient microplate format (Skvortsov, Zvereva, Shpanchenko, & Dontsova, 2011).

Phosphorylated histone H2AX (λ -H2AX) is another sensitive biomarker for *in vitro* analysis of DNA damage by photoaging. H2AX, a component of nuclear histone protein H2A, is rapidly phosphorylated at Ser 139 with the induction of DNA double-strand breaks and plays an important role in the recruitment of DNA repair proteins (Ibuki & Toyooka, 2015). Immunofluorescence staining of λ -H2AX in human keratinocytes have produced lower limits of detection compared to when cell viability and comet assays are used. In addition, λ-H2AX provides for easy antibody detection. As such, λ -H2AX is a sensitive and efficient tool for monitoring UV-induced DNA damage (Cui et al., 2020; Nagelkerke & Span, 2016; Toyooka, Ishihama, & Ibuki, 2011; Valdiglesias, Giunta, Fenech, Neri, & Bonassi, 2013).

## MATERIALS AND METHODS

### Sample Preparation

3 dry commercial *R. rosea* extracts were obtained from different suppliers, i.e., Sample 1 (NOW FOODS, USA), Sample 2 (ROCKER M, Taiwan), and Sample 3 (HOLY, Taiwan) through PChome, a publicly accessible online shopping market. All three products met the selection criteria of being top sellers and were manufactured as powder in soft gelatin capsules of the same size. Sample solutions for the extracts were prepared by dissolving 2 capsules in 10 mL of 70% EtOH, in compliance with the recommended daily usage of *R. rosea*. Each capsule contained approximately 500 mg of *R. rosea* powder; the exact mass of the ingredients in the prepared solutions were 1.2081g, 1.0316g, and 0.9712g for Products 1, 2, and 3, respectively. Undissolved matter was separated by centrifugation.

For comparison, unprocessed natural *R. rosea* roots and rhizomes cultivated in the Tibetan highlands were purchased from a CHM clinic and their contents extracted (Sample 4). The herbal raw materials were finely comminuted then moistened with pure EtOH overnight. The mixture was then purified by centrifugation and filtration. EtOH solvent was removed by rotary evaporation (IKA-Werke, Germany) and the dried powder was dissolved in 70% EtOH (1g/10mL) (for details see Supplementary Data Table 1).

### Reference Standards

Salidroside and rosavin reference standards were purchased from Sigma Aldrich (≥95% and ≥98% purity, respectively) and dissolved in 70% EtOH to create 50mM stock solutions.

### HaCaT Cell Cultures

Human immortalized keratinocyte cell lines (HaCaT) were cultured and subjected to drug and UV treatment. HaCaT cells were maintained in Dulbecco’s modified Eagle’s medium/Ham’s Nutrient Mixture F-12 (DMEM/F-12) with 10% fetal bovine serum and 1% antibiotics (10,000 *μ*g/mL streptomycin and 10,000 units/mL penicillin) at 37°C in a humidified 5% CO2 incubator. Cells were serially passaged at 80-90% confluence. Prior to drug treatment, HaCaT cells were grown to 100% confluence.

### Drug Treatment and UV Irradiation

Cultured HaCaT cells were seeded in 6-well microplates at a concentration of 2 * 10^5^ cells per well. Cells were treated with either 300 μL of extract solution (Samples 1-4) diluted in PBS (1:100) to 200μg/mL, or 900 μL of Salidroside and Rosavin standard stock solution diluted in PBS to 50μg/mL. A vehicle control of 70% EtOH was used. Following drug treatment, cells were incubated at 37°C for 1 hour.

Drug-treated HaCaT cells were then exposed to UV radiation to simulate photoaging. Cells were irradiated with Dermapal phototherapy devices (Daavlin, USA) equipped with a UVA power output of 21.4 mW/cm^2^, and a UVB power output of 7.6 mW/cm^2^. The ideal duration for UVA and UVB irradiation was determined by incrementally increasing exposure time from the minimal erythema dose until visible morphological change was induced. Exposure time was optimized at 4 minutes and 10 seconds for UVB wavelengths, amounting to a total of 2000 mJ. For the weaker UVA wavelengths, exposure time was optimized at 10 minutes and 52 seconds, for a total of 15 J.

### Cell Morphology Assessment

Cell morphology of HaCaT cells was studied as a qualitative biomarker of cell aging. Following drug treatment and UV irradiation, cell morphological changes, such as shape, roughness, volume, as well as cell-cell adhesion were examined using an inverted light microscope (MOTIC AE2000). Images of magnification 200X were taken. Healthier cells displayed normal cell adhesion, while aged cells demonstrated clumping up and a loss of structure.

The anti-aging effects of different extracts and standards were compared by tallying the number of apoptotic cells for 10 randomly chosen magnified fields for each sample. The number of senescent cells between the control group and the samples were compared by means of a t-test. SPSS for windows, version 22.0 (IBM, Armonk, New York), was used to perform statistical analyses, and P values < 0.05 were considered statistically significant.

### Immunofluorescence Staining and Confocal Laser Scanning Microscopy

Keratinocytes were grown on glass coverslips to 90% confluence and subjected to drug treatment and UV irradiation. Cells were then fixed in 4% paraformaldehyde at 4 °C for 10 min and washed three times with 2% bovine serum albumin in PBS for 10 min each. 0.5% Triton X-100 was added for 10 min at 4 °C to induce permeabilization, followed by three 5-min washes, and blocking with 5% goat serum in PBS for 1 hour.

Cells were probed with mouse anti-phospho-Histone H2A.X (Ser139) antibody (1:1000; Sigma-Aldrich) and rabbit anti-β-Actin (1:2000; Sigma-Aldrich) for 2 hours. CF-488A conjugated goat anti-mouse (1:1000; Sigma-Aldrich) and CF-640R conjugated goat anti-human (1:1000; Sigma-Aldrich) secondary antibodies were used to probe for phospho-Histone H2A.X and β-Actin respectively. Coverslips were mounted on slides with proLong gold antifade reagent with 4’,6-diamidino-2-phenylindole (Invitrogen, USA), which was used to probe for the nucleus.

Images were analyzed with a Leica TCS SP8X confocal microscope using excitation wavelengths of 405, 490, and 642 nm. Images were viewed with a 63 objective with a numerical aperture of 1.4. Triple-labeled samples were checked for bleed-through by turning off the respective lasers and assaying for the absence of image. Independent representative images were assembled using Microsoft PowerPoint; brightness and contrast were uniformly adjusted across all images.

### Detection of Telomerase Activity

The *Telo* TAGGG Telomerase PCR ELISA kit (Roche, Basel, Switzerland) was used to detect the telomerase activity of keratinocytes after UV irradiation. The immunoassay consists of two parts: amplification by polymerase chain reaction (PCR) and detection by enzyme-linked immunosorbent assay (ELISA). For each sample, 2 * 10^5^ cells were harvested and lysed, and cellular contents were collected via differential centrifugation.

Prior to amplification, telomerase adds telomeric repeats (TTAGGG) to the 3’-end of the biotin-labeled synthetic P1-TS primer. These elongation products, as well as provided Internal Standards (IS) were added to the same reaction vessel and amplified by PCR using primers P1-TS and the anchor-primer P2. For each sample, negative controls were created by heat treatment of the cell extracts for 10 minutes at 85°C prior to PCR reaction, which inactivated the telomerase protein. The PCR products derived from telomerase-mediated elongation contained telomerase-specific 6 nucleotide increments, while the IS generated a 216 bp product. The cycling process was performed in a GeneATLAS G02 Thermal Cycler (Astec Bio, Japan).

PCR products were then split into two aliquots, denatured, and hybridized separately to digoxigenin-(DIG)-labeled detection probes, specific either for the telomeric repeats (P3-T) or the IS (P3-Std), respectively. The resulting products were immobilized via the biotin label to a streptavidin-coated microplate. Immobilized amplicons were then detected with an antibody against digoxigenin conjugated to horseradish peroxidase (Anti-DIG-HRP), and the colorimetric peroxidase substrate TMB. Color development was allowed to occur for no longer than 5 minutes.

An ELISA plate reader (Tecan, Switzerland) was used to measure the absorbance of the samples at 450 nm, with a reference wavelength of 690 nm, within 30 minutes after adding the stop reagent. Absorbance values were reported as the A450 nm reading against a blank (reference wavelength A690 nm). The quantification of relative telomerase activity was determined by comparing the signal of the samples to the signal obtained from a known amount of Control template (TS8), provided as a ready-to-use solution at a concentration of 0.1 attomol/mL. Results from three independent experiments were expressed as mean +- S.E. Where indicated, data were compared using the Student’s t test. Statistical significance was defined as p < 0.05.

### Analysis of Active Compound

Isolation and quantification of the active compounds in the commercial *R. rosea* samples was achieved with two chromatographic techniques – Prep TLC followed by IR spectroscopy, as well as HPLC. Isolation of rosavin was best achieved used plate-based methods, while isolation of salidroside was best achieved through liquid chromatography.

#### Prep TLC method

Analysis of rosavin was performed using glass-backed preparative thin layer chromatography (Prep TLC) (SILICYCLE, SiliaPlate, 60A). The solvent system used was ethyl acetate-methanol-water (77:13:10, v/v/v). Standard amounts of 0.25 mL *R. rosea* solution was spotted onto the plate. Plates were dried and visualized in a UV viewing cabinet (Newsoft Technology Corporation 42’’ industrial grade display) at 254 nm where the retention factor (rf) of each band was measured. Bands with rf values corresponding to that of reference standards were recovered by scraping the adsorbent layer in the region of interest. The isolated materials were dissolved and eluted in 95% EtOH, filtered, then run through a rotary evaporator (RV 10; IKA, Staufen, Germany) for solvent removal. The dried residue was collected for analysis.

An IR spectrometer (PerkinElmer UATR Two) was used to confirm the identity of the active components collected. The IR spectra of the collected residues were compared to the IR spectra of the reference standards for matching peaks. Once their identity was confirmed, the weight percent of each active component in the commercial extracts was determined by dividing the mass of the isolated compound by the total mass of the spotted sample.

#### HPLC method

Isolation and quantification of salidroside was achieved using high-performance liquid chromatography (HPLC). Samples were separated on an InertSustain C18 column (4.6 × 250 mm, 5μm) (GL Sciences, Japan) with a mobile phase of 13% methanol at a flow rate of 1.0 mL/min. The chromatograms were registered at 203 nm. The injection volume was 20 μL.

## RESULTS

UV irradiated keratinocytes were pretreated with *R. rosea* extracts and reference standards then examined for aging biomarkers both qualitatively and quantitatively, using light microscopy, confocal microscopy, and TRAP-ELISA. Data obtained from these tests were used to validate and compare the anti-aging effects of the *R. rosea* samples. In parallel, isolation of rosavin and salidroside active compounds in the extracts was performed with Prep-TLC and HPLC, respectively. Chromatography results combined with results from treatment of keratinocytes indicate the ingredients responsible for *R. rosea’s* anti-aging capabilities.

### Quantitative analysis of *R. rosea* treated keratinocytes shows fewer senescent cells and higher telomerase activity

Drug-treated and UV-irradiated cells captured by images taken at 200x were examined qualitatively and classified as either healthy or aged cells. For each sample, the number of aged cells for ten randomly chosen fields were recorded and compared by means of a t-test (Table 1). For both UVA and UVB irradiated cells, the aged cell count of *R. rosea* test samples were significantly less than that of the EtOH control. In particular, test samples 1 and 4 had fewer aged cells compared to samples 2 and 3. Salidroside and rosavin treated reference standards were also among the samples with the lowest aged cell count. Samples radiated with UVB produced more aged cells than samples radiated with UVA.

**Table 1.** Average Aged Cell Count for *R. rosea* test samples under UVA and UVB.

|  | UVA |  |  | UVB |  |  |
| --- | --- | --- | --- | --- | --- | --- |
|  | Aged Cell Count / HPF | P-Value | 95% CI | Aged Cell Count / HPF | P-Value | 95% CI |
| **70% EtOH Control** | 15.08 ± 2.74 |  |  | 17.00 ± 3.38 |  |  |
| **Test Sample 1** | 8.29 ± 2.57 | <.001 | 4.89 - 8.68 | 10.79 ± 2.22 | 0.001 | 2.99 - 9.43 |
| **Test Sample 2** | 10.05 ± 2.23 | <.001 | 4.21 - 7.84 | 12.53 ± 3.33 | 0.013 | 1.03 - 7.89 |
| **Test Sample 3** | 10.47 ± 3.79 | 0.002 | 1.92 - 7.31 | 12.60 ± 2.55 | 0.009 | 1.19 - 7.60 |
| **Test Sample 4 (Natural Root)** | 9.10 ± 1.44 | <.001 | 3.96 - 7.99 | 9.88 ± 3.50 | <.001 | 3.72 - 10.52 |
| **Salidroside** | 8.10 ± 1.86 | <.001 | 5.32 - 8.64 | 8.62 ± 1.93 | <.001 | 5.10 - 11.66 |
| **Rosavin** | 8.42 ± 2.65 | <.001 | 4.83 - 8.48 | 9.31 ± 2.05 | <.001 | 4.39 - 10.99 |

Telomerase activity of the samples was detected by TRAP-ELISA assay. Relative telomerase activity units (RTA) were calculated according to the following formula:

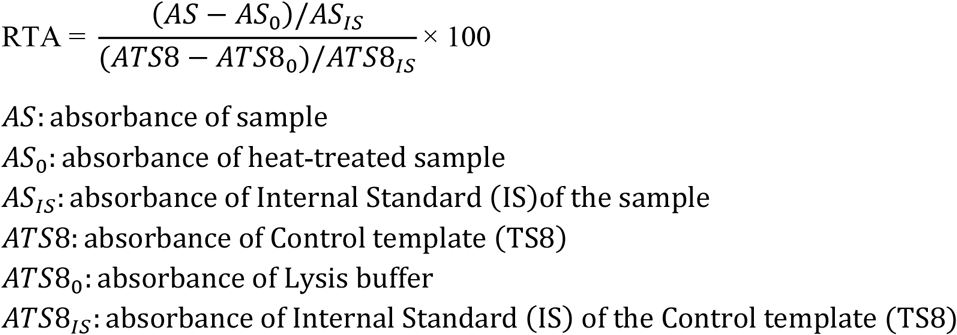

The RTA of UVA and UVB irradiated cell samples were plotted as bar graphs with ± 1 S.E. (Fig. 1). For both UVA and UVB wavelengths, cells treated with *R. rosea* extracts and cells treated with salidroside and rosavin reference standards displayed a significantly higher RTA than that of the EtOH control. For both wavelengths, Sample 1 shows a higher RTA value than the other commercial extracts. Sample 4, treated with natural *R. rosea* root, also displayed high telomerase activity. While the RTA of salidroside and rosavin samples are similar under UVB, the RTA of salidroside is significantly higher than rosavin under UVA.

**FIGURE 1.**
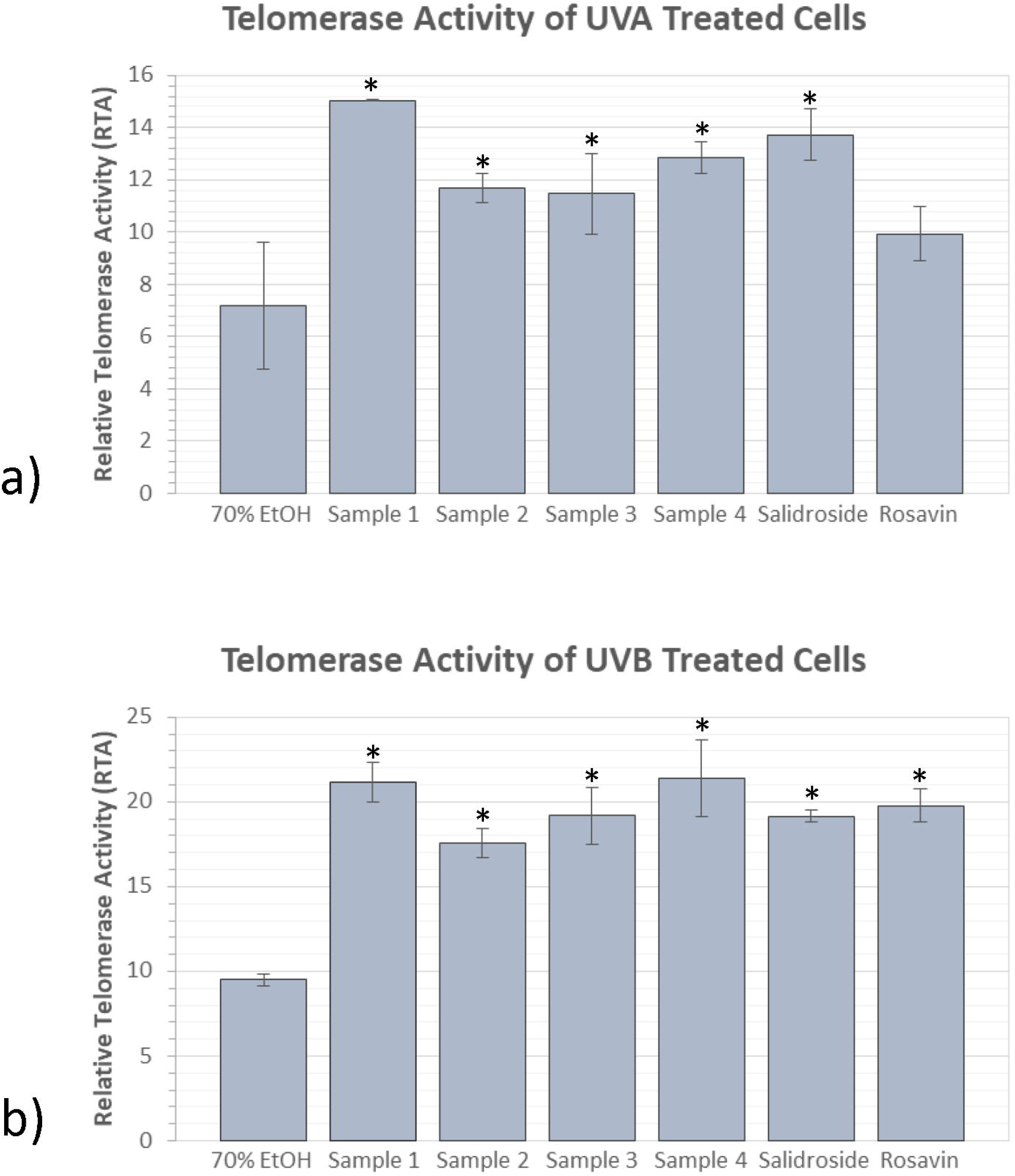
Bar graphs shows the relative telomerase activities of UV irradiated cells treated with commercial extracts, natural root, reference standards, and 70% EtOH. Error bars of ± 1 S.E are shown. Cells were exposed to a) 15J of UVA and b) 2000 mJ of UVB. *R. rosea* treated samples had significantly higher telomerase activity than that of the control. Sample 1 in particular had the greatest telomerase activity.

### Detection of γ-H2A.X positive cells by confocal laser scanning microscopy yields results in agreement with quantitative analysis

Figure 2 shows the generation of γ-H2A.X in cultured HaCaT cells following irradiation with both UVB (2000mJ) and UVA (15J). γ-H2A.X foci within the nucleus (co-localization with blue DAPI) were clearly detected as green dots. The number of cells with more than 10 foci was significant decreased by pre-treatment with *R. rosea* extracts, as well as salidroside and rosavin. Moreover, images magnified to depict single cells show extensive homogenous staining of γ -H2A.X in control cells. Contrastingly, the γ-H2A.X staining in *R. rosea* treated cells appeared as discrete dots with less density. For both UVA and UVB, Sample 1 showed much less γ-H2A.X positive cells than other commercial extracts. Cell treated with sample 4 (natural *R. rosea* root) also displayed strong protective effects. Interestingly, after UVA irradiation, salidroside but not rosavin treated cells had reduction in γ -H2A.X generation.

**FIGURE 2.**
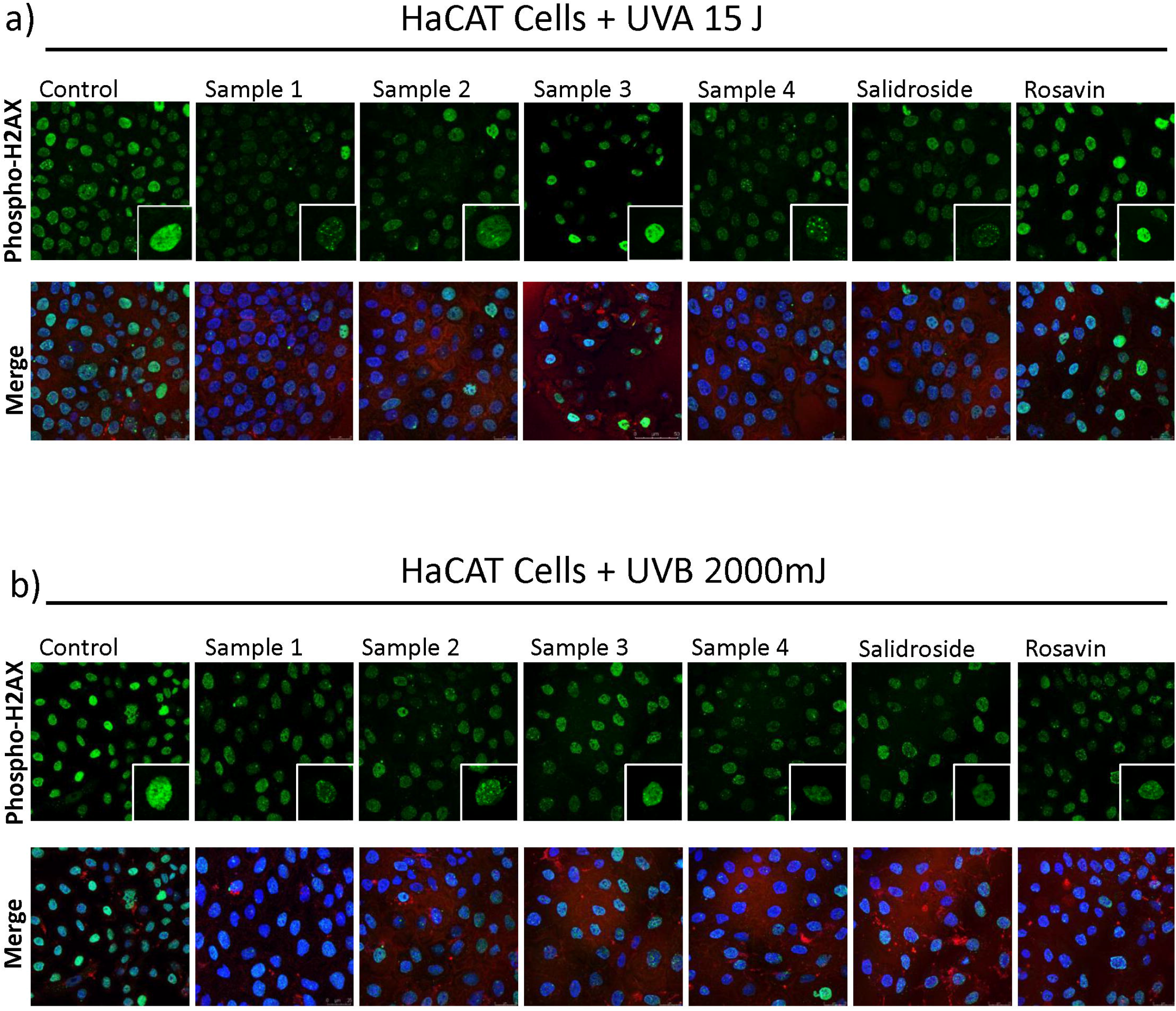
Drug-treated and UV-irradiated HaCaT cells were stained with (i) anti-phospho-Histone H2A.X, a marker of DNA damage (green), (ii) anti-β-Actin (red) for cytoskeleton, and (iii) DAPI for nuclear localization (blue). Cells were irradiated with both a) UVA (15J) and b) UVB (2000mJ). R. rosea treated cells had less γ-H2A.X foci and more dispersed staining compared to that of the control.

### Commercial *R. rosea* extracts vary considerably in salidroside and rosavin content

TLC of *R. rosea* extracts was tested on various solvent systems consisting of water, methanol, ethyl acetate, hexane, acetone, and acetic acid. Ethyl acetate-methanol-water (77:13:10, v/v/v) was selected for the mobile phase as it yielded distinct bands corresponding to salidroside and rosavin when simultaneous TLC was performed (Fig. 3). The same mobile phase was then used to perform Prep TLC, where *R*_*F*_ values of the reference standards were used to locate bands of interest. These bands were isolated and purified, and their identity confirmed by IR Spectroscopy (Fig. 4). By dividing the mass of the purified band by the total mass of the applied sample, we determined the weight percent of the compounds. Successful isolation and quantification of rosavin, but not salidroside was achieved in all 3 *R. rosea* samples using this method. Isolation of salidroside was achieved using HPLC (Fig. 5). To determine the weight percent of salidroside from the chromatograms, the known concentration of salidroside standard was multiplied by the ratio of compound peak area to standard peak area. The weight percent of salidroside and rosavin in the 3 commercial *R. rosea* samples are shown in Figure 6. Sample 1 contained the greatest concentration of both salidroside and rosavin, with 13.70% rosavin and 7.74% salidroside. Samples 2-3 contained similar concentrations of active ingredients, significantly less than that found in Sample 1. There is still a large percentage of compounds present in the commercial *R. rosea* samples that have yet to be isolated and analyzed.

**FIGURE 3.**
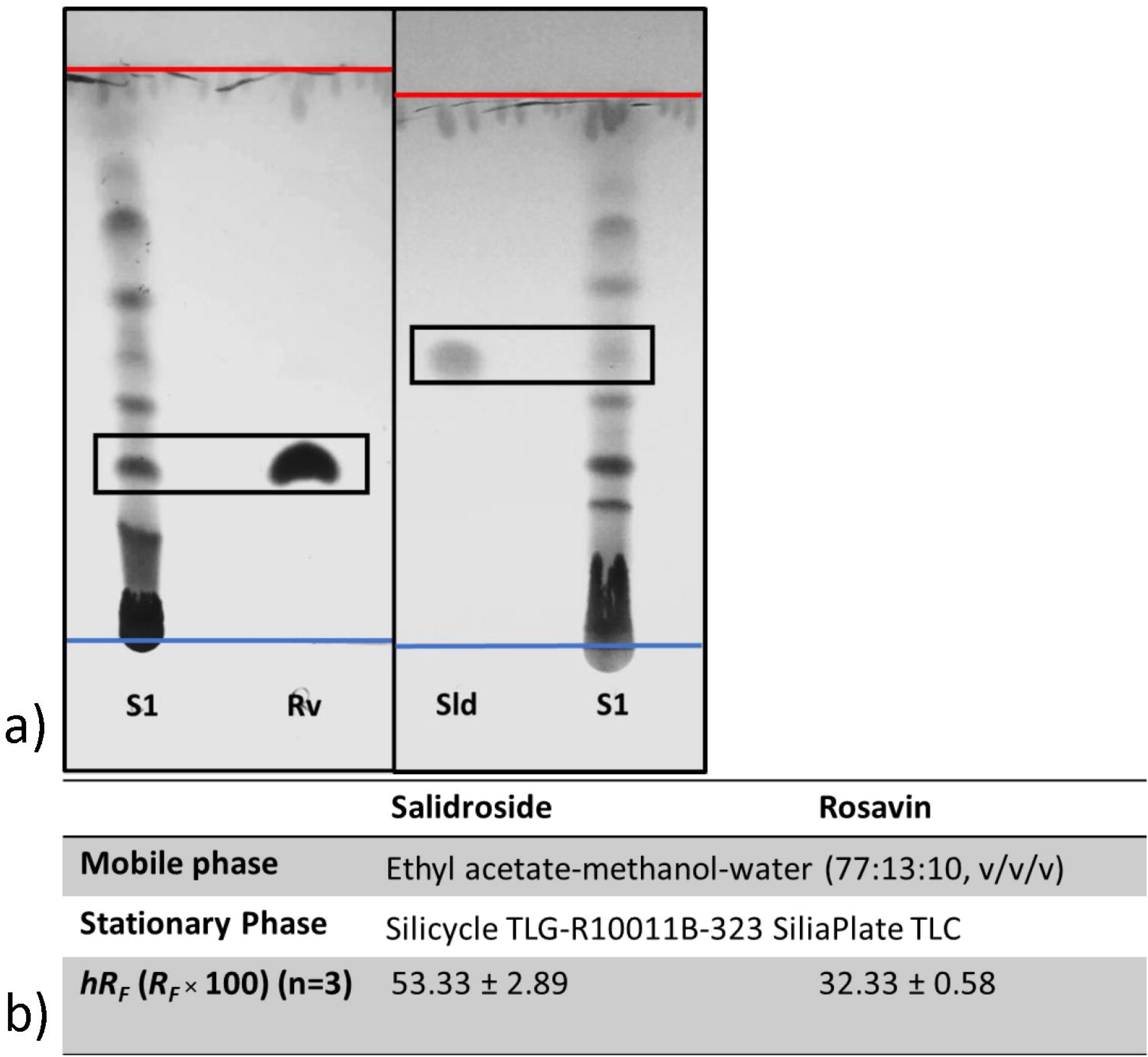
Simultaneous TLC of commercial *R. rosea* extract (S1) and reference standards (Rv and Sld). a) Plates were visualized at 254 nm and images were adjusted to maximize “contrast” and “pop”. The blue line marks the origin, while the red line marks the solvent front. b) *R*_*F*_ values for salidroside and rosavin were measured and averaged across 3 trials.

**FIGURE 4.**
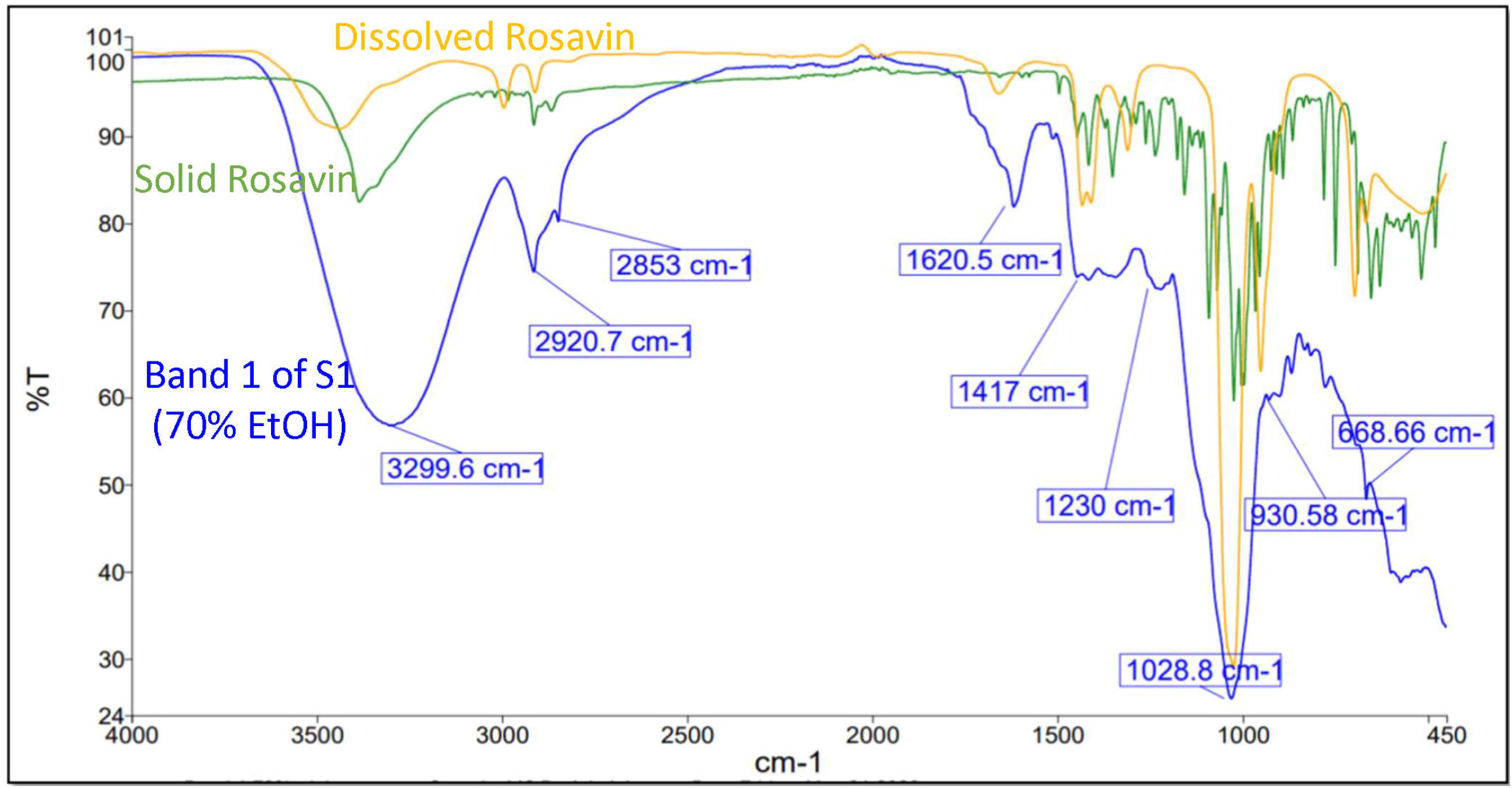
Comparison of the IR spectrum of isolated band (S1) with the spectrum of rosavin reference standards (in solid and dissolved states). Matching peaks at 3299.6 cm^-1^, 2920.7 cm^-1^, 2853 cm^-1^, etc. indicate successful isolation of rosavin in Sample 1. This same process was then repeated to isolate rosavin in Samples 2 and 3.

**FIGURE 5.**
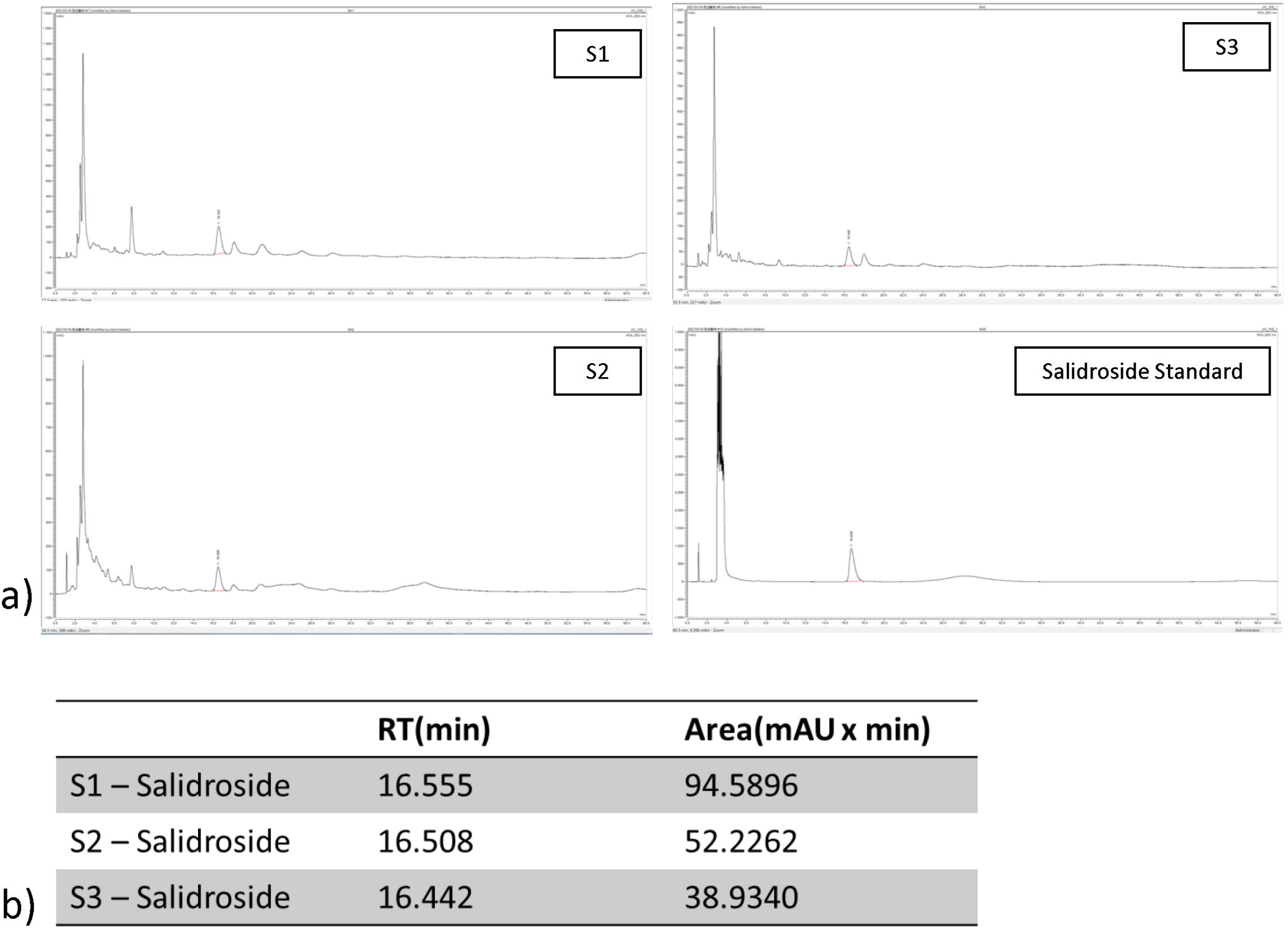
a) HPLC chromatograms of *R. rosea* extracts (S1 – S3) and salidroside reference standard. b) Quantitative data for salidroside peaks in the extracts.

**FIGURE 6.**
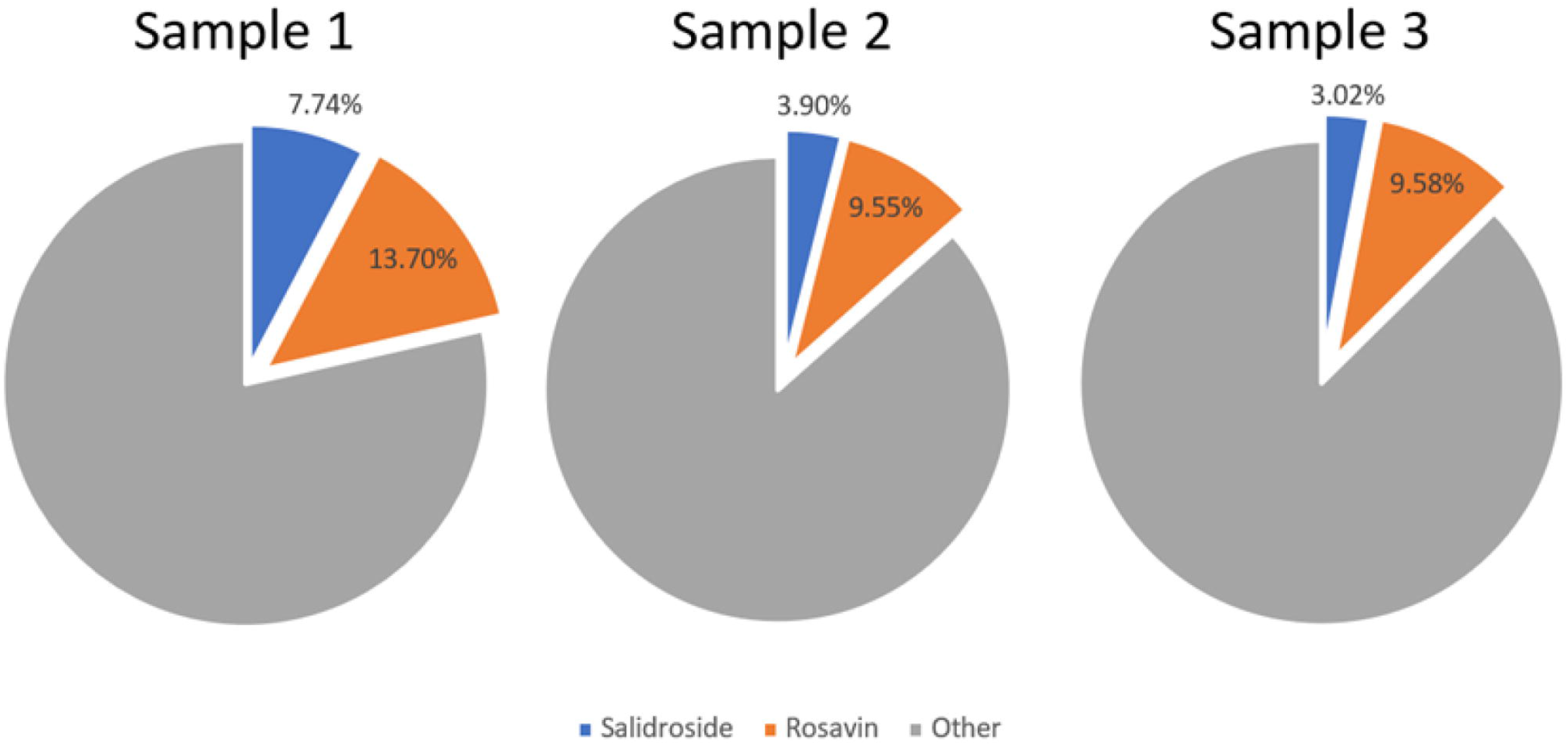
Pie chart depicts the composition of active ingredients in the 3 commercialized *R. rosea* samples. The weight percent of rosavin was found to be 13.7% in Sample 1, 9.55% in Sample 2, and 9.58% in Sample 3. The weight percent of salidroside was found 13.70% in Sample 1, 3.90% in Sample 2, and 3.02% in Sample 3. The gray portion of the pie chart represents compounds that have yet to be isolated and identified.

## DISCUSSION

In recent years, the increasing use of CHM has popularized anti-aging herbs like *R. rosea*. But despite this recognition, CHM’s acceptance into global healthcare is hindered by a lack of standardization of ingredients, as well as insufficient knowledge of the herbal medicine’s anti-aging ability. In this study, the evaluation of biological markers of aging produced largely corroborative results. *R. rosea* treated cells had less γ-H2A.X staining, higher telomerase activity, and lower aged cell counts compared that of the control. As such, morphological and quantitative analysis of *R. rosea’s* anti-aging effects both confirm *R. rosea’s* ability to rescue UVA and UVB induced changes in keratinocytes.

*R. rosea’s* preventative effects, however, are not uniform across commercial *R. rosea* samples. While all *R. rosea* samples had significantly greater preventative effects that than that of the control, certain samples displayed greater efficacy than others. Analysis of the active ingredients in the *R. rosea* samples reveals not only that the samples vary in composition, but that samples with greater salidroside and rosavin content possess significantly greater anti-aging effects. It is thus noted that the concentration of active ingredients in *R. rosea* extracts play a key role in determining the extent of the product’s preventative effects. This observation further emphasizes the importance of standardization in CHM.

The results presented in this study also provide insight on the exact nature of *R. rosea’s* anti-aging effects. The telomerase activity of drug treated keratinocytes differ under UVA and UVB exposure. Specifically, rosavin treated cells had a significantly greater telomerase activity than the control when irradiated with UVB but not UVA. This characteristic was confirmed by confocal microscopy, where a reduction in γ-H2A.X generation was not observed in rosavin treated samples under UVA. The different wavelengths that characterize UVA and UVB light produce different degrees of penetration and intensity. It is not unlikely that rosavin’s preventive effects are most prominent only within specific ranges of light. Histological studies have found that while UVB is responsible skin cancer, UVA irradiation markedly induces MMP activity, enhancing the degradation of collagen, leading to apoptosis of fibroblasts and the evocation of inflammatory cells (Wang et al., 2019). Full coverage of the UV spectrum is therefore crucial and may necessitate *R. rosea* products that contain other active compounds in addition to rosavin.

It is also noted that certain *R. rosea* samples, namely Sample 1, exhibit greater preventative effects than both salidroside and rosavin standards. This points to the potential existence of synergistic interactions between components in *R. rosea*, where complex mechanisms generate a combined effect greater than the sum of individual effects. This property of CHM has long been speculated but evidence to support this theory remains inadequate (Zhou et al., 2016).

Overall, this study affirms *R. rosea’s* potential as an anti-aging herbal remedy for the skin. Such a cosmetic product is sure to have immense market value. However, several steps must still be taken if *R. rosea’s* potential is to be realized. First, more comprehensive analysis of the composition of *R. rosea* extracts must be performed. While this study reports the quantification of salidroside and rosavin, the two most frequently discussed compounds in literature, it is likely there are other phenolic compounds present in *R. rosea* samples that contribute to their anti-aging effects. Understanding the role these compounds play is crucial to the proper integration of modern scientific techniques with traditional knowledge. Furthermore, the precise relationship between the concentration of phenolic compounds and corresponding effects can be analyzed with the development of a dose-response curve. This will allow for the identification of ideal ingredient concentrations to maximize preventative effects.

*R. rosea’s* incorporation into a feasible cosmetic product will require further experimentation with product stability and efficient mechanisms of delivery. Nonetheless, the findings in this study reaffirm *R. rosea’s* efficacy as a pharmacological anti-aging remedy and provide a basis for CHM’s integration into the mainstream of global healthcare.

## CONFLICT OF INTEREST

The authors declare that the research was conducted in the absence of any commercial or financial relationships that could be construed as a potential conflict of interest.

## DATA AVAILABILITY STATEMENT

The datasets generated for this study are available on request to the corresponding author.

## AUTHOR CONTRIBUTIONS

WHT and HEL conceptualized the study. WHT performed the experiments. WHT, CBC, JC, and CCH analyzed the data. JC, HEL, and WHC supervised the study. WHT wrote the first draft of the manuscript. All authors contributed to the article and approved the submitted version.

## FUNDING

This work was supported in part by grants from Chang Gung Memorial Hospital (CMRPG1F0072).

## ACKNOWLEDGEMENTS

The authors would like to thank the faculty of microscopy core laboratory of Chang Gung Memorial Hospital with technical assistance in confocal imaging. The technical expertise of Vercotech Inc. in the area of HPLC is also deeply appreciated. Lasty, the authors thank the Science Department and Sean Tsao of Taipei American School for guidance and resources.

## Supplementary Data

**Table S1.**
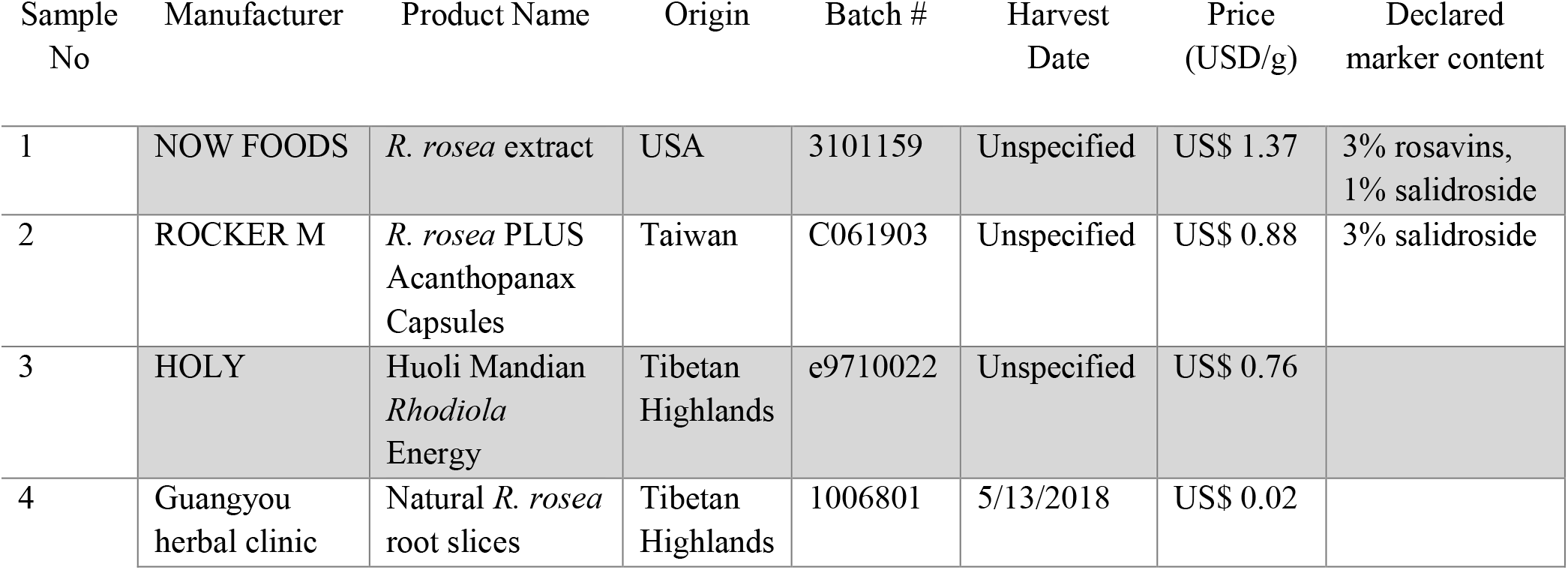
Test samples’ manufacturer, origin, batch number, harvest date, price per gram, and declared marker content.

| Sample No | Manufacturer | Product Name | Origin | Batch # | Harvest Date | Price (USD/g) | Declared marker content |
| --- | --- | --- | --- | --- | --- | --- | --- |
| 1 | NOW FOODS | *R. rosea* extract | USA | 3101159 | Unspecified | US\$ 1.37 | 3% rosavins, 1% salidroside |
| 2 | ROCKER M | *R. rosea* PLUS Acanthopanax Capsules | Taiwan | C061903 | Unspecified | US\$ 0.88 | 3% salidroside |
| 3 | HOLY | Huoli Mandian *Rhodiola* Energy | Tibetan Highlands | e9710022 | Unspecified | US\$ 0.76 | |
| 4 | Guangyou herbal clinic | Natural *R. rosea* root slices | Tibetan Highlands | 1006801 | 5/13/2018 | US\$ 0.02 | |

## Notes

### Competing Interest Statement

The authors have declared no competing interest.

